# Hi-C deconvolution of a textile-dye degrader microbiome reveals novel taxonomic landscapes and link phenotypic potential to individual genomes

**DOI:** 10.1101/2020.06.18.159848

**Authors:** Ayixon Sánchez-Reyes, Luz Bretón-Deval, Hayley Mangelson, Ilse Salinas-Peralta, Alejandro Sanchez-Flores

**Affiliations:** Cátedras Conacyt-Instituto de Biotecnología, Universidad Nacional Autónoma de México; Phase Genomics Inc.; Universidad Politécnica del Estado de Morelos; Instituto de Biotecnología, Universidad Nacional Autónoma de México

**Keywords:** Hi-C metagenomic deconvolution, textile-dye biodegradation, metagenome assembled genomes

## Abstract

Microbial biodiversity is represented by genomic landscapes populating dissimilar environments on earth. These genomic landscapes usually contain microbial functional signatures connected with the community phenotypes. Here we assess the genomic microbiodiversity landscape of a river associated microbiome enriched with 200 mg.mL^−1^ of anthraquinone Deep-Blue 35 (™); we subjected to nutritional selection a composite sample from four different sites from a local river basin (Morelos, Mexico). This paper explores the resultant textile-dye microbiome, and infer links between predicted biodegradative functions and the individual genome fractions. By using a proximity-ligation deconvolution method, we deconvoluted 97 genome composites, with 80% of this been potentially novel species associated with the textile-dye environment. The main determinants of taxonomic composition were the genera *Methanobacterium*, *Clostridium*, and *Cupriavidus* constituting 50, 22, and 11 % of the total population profile respectively; also we observe an extended distribution of novel taxa without clear taxonomic standing. Removal of 50% chemical oxygen demand (COD) with 23% decolorization was observed after 30 days after dye enrichment. By metagenome wide analysis we postulate that sequence elements related to catalase-peroxidase, polyphenol oxidase, and laccase enzymes may be causally associated with the textile-dye degradation phenotype under our study conditions. This study prompts rapid genomic screening in order to select statistically represented functional features, reducing costs, and experimental efforts. As well as predicting phenotypes within complex communities under environmental pressures.

## 1. Introduction

Metagenomics approaches have been largely applied to characterize microbial communities’ structure and functions in different environments. Two major goals on metagenomic analysis of complex communities are to profile and compare the taxonomic composition and infer their associated phenotypic potential; in order to get a comprehensive understanding of environmental microbiodiversity and its concomitant effects. Recently, the recovering of individual genomic complements from metagenomes (MAGs) has become an attractive task, generating new insights about uncultured specimens usually underrepresented in customs genome data repositories and discovering totally new microbial genomes-landscapes [1].

Historically, the ecosystems most scrutinized by metagenomic have been host-associated or environmental origin [2, 3]. Among the aquatic ecosystems, two of the most neglected biomes involve freshwater and sediments metagenomes, despite the importance of microbial dynamics in the function of freshwater surface bodies [4]. Since 2010, 10,648 metagenomes have been uploaded to the National Center for Biotechnology Information (NCBI) Assembly database, 148 from freshwater environments, and 111 from sediments (https://www.ncbi.nlm.nih.gov/assembly/?term=derived+from+metagenome consulted 21/03/2020). Arguably, only a few hundred of individual genomes have been assembled from these metagenomes (https://www.ncbi.nlm.nih.gov/assembly/?term=metabat consulted 21/03/2020). Furthermore, the Integrated Microbial Genomes & Microbiomes (IMG/M) system contains 9898 metagenomes, from which 13,866 bins from freshwater environments have been assembled (761 bins from river samples and 508 from sediments), representing just 1.58% of total metagenomes bins (https://img.jgi.doe.gov/cgi-bin/m/main.cgi?section=MetagenomeBins&page=bins&type=ecosystem consulted 21/03/2020).

Often it is difficult to connect the ecological functions that take place in microbiomes with the specific biotypes that perform them. Binning methods have brought us closer to inferring individuals that might be performing discrete functions within a mixed sample. The binning methods offer an attractive and committed way to assemble genomes from metagenomes [5, 6]. However, these methods are not without limitations *v. gr.*, low reconstruction genome rate, or bins clustering with low quality, usually associated with the incorporation of spurious reads and the dependence on two-dimensional parameters like compositional regularity of bases among others [7]. Genome-resolved metagenomics still struggles to achieve high quality and contiguity indicators.

Recently, methods based on chromosomal proximity ligation (Hi-C or 3C) have been applied to extract genomes from metagenomes with reasonable indicators of quality, taking physical neighborhood signals of sequences that are co-located in cells [8, 9]. Several papers show their applicability to deconvolve microbiomes and extract functional-phenotypic determinants related to community performance [10–12].

To assess the genomic microbiodiversity landscape that can be resolved using proximity ligation (Hi-C) chemistry in a river associated microbiome, we subjected to nutritional selection - with a proprietary oil blue family anthraquinonic dye-a composite sample from sediment and freshwater from Apatlaco river basin (Morelos, Mexico) for 30 days in batch conditions, to enrich a textile-dye degrader community fraction. From there, we deconvolved 97 genome composites and evaluated the overall dye-removal achieved by the mixed culture. This paper explores the resultant textile-dye microbiome (TDM), in order to infer links between predicted biodegradative functions and the individual genomic complements of the community. We deal with the hypothesis that it is possible to connect which individual genomic composites may contribute a given function within a community under nutritional selection. This is the first report about taxa-specific Hi-C deconvolution analysis in a textile dye related biome, and the first approach to deconvolve the Apatlaco River microbiome, the basin most impacted by human activity in the state of Morelos.

## 2. Material and Methods

### 2.1. Sample recovery, DNA preprocessing and Hi-C deconvolution analysis

The Apatlaco river basin (Morelos, México) was our study model to explore the effects of anthropogenic pollution on microbial diversity. Four samples of sediment and surface water were taken and processed as described by Breton-Deval *et al*., [5] (sites S1: −99.26872, 18.97372, S2: −99.2187, 18.83, S3: −99.23337, 18.78971 and S4: −99.18278, 18.60914). One composite sample (~ 2 L) was enriched in the laboratory with 200 mg. mL^−1^ of Deep-Blue 35(™), an anthraquinone dye from the oil blue family as sole carbon source. The enriched sample was incubated for 30 days at room temperature in a 10 L batch reactor. Ten grams of the sediment sludge plus 10 mL of water column were extracted from the reactor and directly cross-linked according to [11]. A Hi-C library was created from the sample using a Phase Genomics (Seattle, WA) ProxiMeta Hi-C Microbiome Kit, which is a commercially available version of the Hi-C protocol [13]. Following the manufacturer’s instructions for the kit, intact cells from the sample were simultaneously digested using the *Sau3A*I and *Mluc*I restriction enzymes, and proximity ligated with biotinylated nucleotides to create chimeric molecules composed of fragments from different regions of genomes that were physically proximal *in vivo*. Continuing with the manufacturer’s protocol, molecules were pulled down with streptavidin beads and processed into an Illumina-compatible sequencing library. Sequencing was performed on an Illumina HiSeq4000, generating a total of 72,007,031 Hi-C read pairs. All metagenomic sequencing files were uploaded to the Phase Genomics cloud-based bioinformatics portal for subsequent analysis.

Previous shotgun reads obtained by Breton-Deval et al., [4] were combined with Hi-C reads and were filtered and trimmed for quality, then normalized using bbtools [14] and assembled with MEGAHIT [15] using default options. Hi-C reads were then aligned to the assembly following the Hi-C kit manufacturer’s recommendations. Briefly, reads were aligned using BWA-MEM [16] with the −5SP and -t 8 options specified, and all other options default. SAMBLASTER [17] was used to flag PCR duplicates, which were later excluded from the analysis. Alignments were then filtered with samtools [18] using the -F 2304 filtering flag to remove non-primary and secondary alignments. Metagenome deconvolution was performed with ProxiMeta [11]. Clusters were assessed for quality using CheckM [19] and assigned preliminary taxonomic classifications with Mash [20]. Binning refiner and DAS tool were used to select optimized and non-redundant set of metagenomes assembled genomes from the initial 97 clusters [7, 21].

### 2.2. Community taxonomic profiling

We estimated microbial community composition by mapping the Hi-C reads against the representative clade-specific markers catalog contained on the MetaPhlAn tool [22]. The abundance profiles obtained were merged and processed with GraPhlAn to producing an abundance-based cladogram tree representation [23]. MetaPhlAn raw abundance tables were visually explored with excel spreadsheet developed by Microsoft in order to identify patterns and trends.

### 2.3. Decolorization of the textile dye Deep-Blue 35(™) by the enriched microbiome

The textile dye Deep-Blue 35(™) was obtained from Monroe Chemical Company de México, S.A. de C.V, in his national commercial form. A dye stock solution (32 g.L^−1^, final pH 9.2) was prepared using deionized water. We evaluated the overall dye-removal achieved by the mixed culture by adding Deep-Blue 35 to the bath reactor up to a final concentration of 200 mg.L ^−1^ as sole carbon source. After 30 days of incubation at room temperature, we assessed the final chemical oxygen demand (COD) using a standard HACH® procedure followed by the spectrophotometric determination in a Hach spectrophotometer DR/4000 UV-Vis. Also, we recorded the UV-Vis spectrum of the sample to evaluate the degree of color removal.

### 2.4. Functional annotation and phenotypic potential inference

The individual genomes composites deconvolved in this study were annotated with Prokka: Rapid Prokaryotic Genome Annotation tool [24]. Besides, they were annotated with KofamKOALA tool to assign KEGG Orthology related to the decolorization of textile dyes [25]. We selected several enzymatic dye-biodegradation markers according to Sarkar *et al*., [26]; after we look for their representation in the resolved genomes and we compared their abundance against previous freshwater metagenomes assembly annotations from [4, 27]. The enzymes selected were: Dye decolorizing peroxidase [EC:1.11.1.19, K15733]; Azoreductase [EC 1.7.1.6, K03206; EC:1.7.1.17, K01118]; Laccase [EC:1.10.3.2; K05909]; Lignin peroxidase [EC:1.11.1.14, K23515]; Tyrosine phenol-lyase [EC:4.1.99.2, K01668]; Tyrosinase [EC:1.14.18.1, K00505]; Catalase-peroxidase [EC:1.11.1.21, K03782]. KEGG mapper was employed to explore pathways and functional modules representative of the textile dye microbiome [28]. Alternatively, we apply the Seer tool [29] in order to investigate sequence signatures (k-mers) significantly associated with the textile dye biodegradation phenotype, assumed from genotype. For this, we associate each deconvoluted genome with the presence or absence for any enzymatic dye-biodegradation marker (already mentioned) in a binary file, were 0 imply no biodegradation phenotype and 1 suppose a potential biodegradative phenotype.

### 2.5. Genus and species boundaries estimation on the genome composites

To assign a taxonomic context for all deconvolved genomes complements of this study, we apply a hybrid approach involving the extraction of rDNA coding sequences from the bins with Barrnap tool when possible and comparing them against homologous sequences from type material [30]. According to this criterion, each genome was placed in the most probable phylogenetic context at the genus or species level. When the rDNA gene resolution was insufficient or absent, Mash distance (D) was estimated against all NCBI type genomes from the most plausible linage for each bin resolved [20]. Type genomes with lower D values were selected for further analysis using average nucleotide identity (ANI) determination. When D ≤ 0.05 or ANI > 95% species context was assigned unambiguously. Novel species were inferred when D > 0.05 or ANI < 95 %. Novel higher taxonomic ranks like genus or family were carefully evaluated by adding up-to-date phylogenomic context comparisons among the most similar type genomes according to Mash [31]. Otherwise, Barco et al., [32] recommendations were set to genus delineation relying on the Alignment Fraction (AF) and ANI values for genus demarcation boundaries (means of 0.331 and 73.98 % respectively). This combined approach ensures unambiguous taxonomic standing assignation for any genome composite moderately complete (> 50 %) and marker gene overrepresentation (< 10 %).

### 2.6. Statistic analysis

Chi-Square exact test was applied to assess KEGG molecular functions (KO) overrepresentation accordingly to functional assignations on each metagenome.

## Data availability

The raw data that support the findings of this study were submitted to NCBI under BioProject accession number: PRJNA623057 and are openly accessible with the following link: https://www.ncbi.nlm.nih.gov/sra/PRJNA623057. Any other data are available from the corresponding author, [ASR], upon reasonable request.

## 3. Results and Discussion

### 3.1. Taxonomic profiles comparison between a textile-enriched and Apatlaco freshwater natural microbiomes

We first characterized the TDM microbiome coming primarily from the Apatlaco river composite samples and compared them with four freshwater microbiomes available from the same basin [4, 27]. The textile dye build-environment involved a considerable abundance of archaea (> 50% pertaining to *Euryarchaeota*, the most abundant phylum). While bacterial phyla *Firmicutes*, *Proteobacteria*, and *Actinobacteria* represented roughly half of the community. MetaPhlAn revealed at least three major clades with abundances >0.5%, containing 10 genera and 14 species. *Methanobacterium* was consistently the most abundant genus; it represented ~50% of the bacterial community in the textile dye microbiome. Also, *Clostridium* (22%) and *Cupriavidus* (~11%) complete the most significant fraction of the metagenome. Overall, the TDM microbiome account for 28 genera (highlighting *Methanobacterium*; *Clostridium*; *Cupriavidus*; *Mesorhizobium*; *Ralstonia*; *Saccharomonospora*; *Sporolactobacillus*); and 41 species-level assignments (highlighting *Methanobacterium paludis*; *Clostridium tyrobutyricum*; *Cupriavidus metallidurans*; *Methanobacterium congolense*; *Mesorhizobium* sp. UASWS1009; *Saccharomonospora azurea*; *Ralstonia* sp. MD27; *Sporolactobacillus laevolacticus*; *Clostridium diolis*) (Figure 1a, Supplementary Material 1). This confirms a diverse community with phylotypes characteristics from oxygen-limited environments and obligate aerobic entities. It is worthy to note that TDM microbiome experienced a drastic reduction on diversity compared with S1-S4 microbiomes [4, 27], sharing only two species-specific lineages: *Variovorax paradoxus* with S1and S4, and *Cutibacterium acnes* with S2 and S4, suggesting a taxonomic context fundamentally distinct from that of natural biomes; maybe due to anthraquinone Deep Blue 35 dye nutritional selection effects (Figure 1b). The co-occurrence of both taxa is not uncommon in the referred conditions. *V. paradoxus* is a highly adaptable microorganism with demonstrated capabilities for anthropogenic contaminants biodegradation [33, 34]. *C. acnes* is a ubiquitous anaerobic *Actinobacteria* part of the human healthy skin microbiota. Metabolically it can degrade triacylglycerol on lipid environments [35]. As *C. acnes* is also found in the human intestine and urinary tract, it is evident a waste-water associated origin, where it could survive colonizing fatty components material.

**Figure 1.**
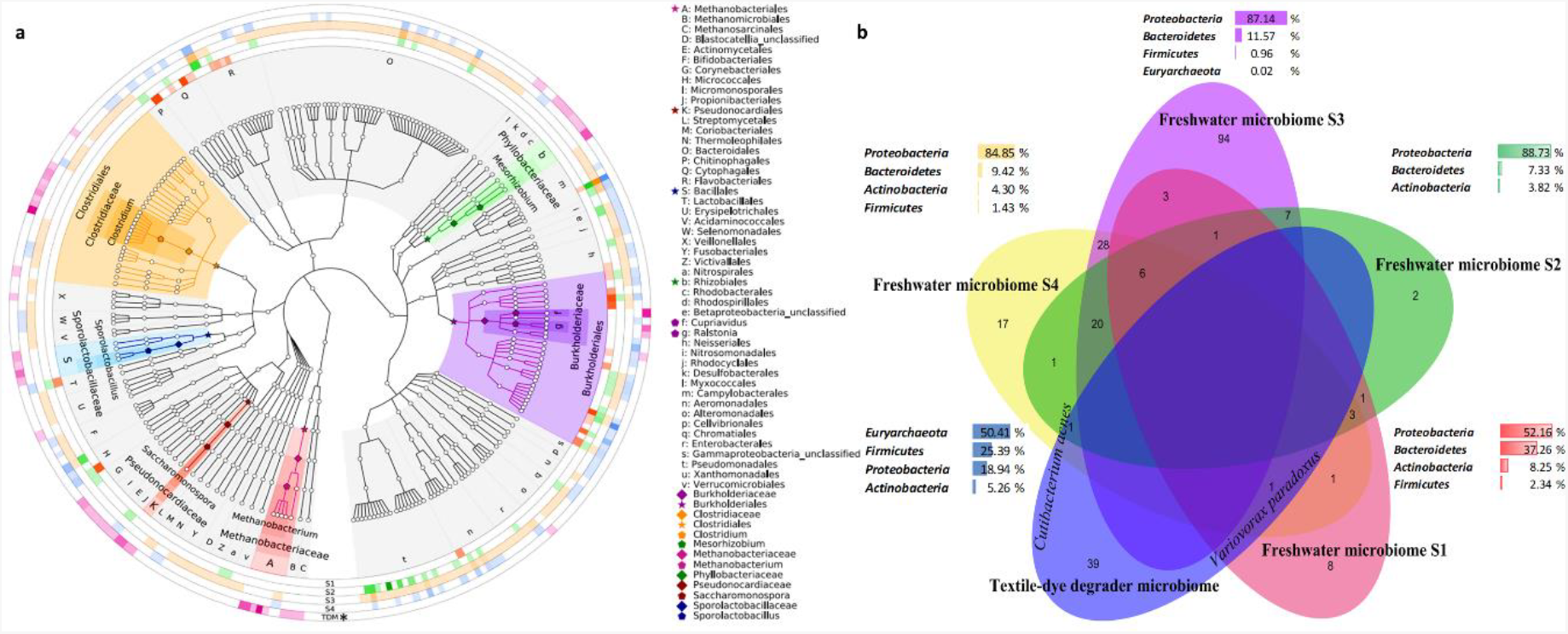
Short-reads metagenomic profiling for the textile-dye selected microbiome (TDM) and four related freshwater microbiomes. GraPhlAn cladogram showing the taxonomic distribution represented as different colors inside the rings. The main determinants in the TDM microbiome are highlighted within the cladogram in different colors and their taxonomic level was identified with text notes (a). Venn diagram showing the distribution at the species level among the compared microbiomes and their composition at the Phylum level (b).

Since the species coverage depends on sequenced microorganisms, MetaPhlAn species-level detection should be taken with caution in light to offer further details on the structure of this metagenome. However, in TDM seems to be apparent the prevalence of methanogenic phylotypes (*Methanomada*, *Stenosarchaea*, and Clostridia groups;) that populate the microbiome and could be important in carbon and CH_4_ cycles.

*Methanobacteriales* and *Clostridiales* (accounting for > 70% of relative abundance), both orders known to contain anoxygenic-types, suggesting a discrete prevalence of anaerobic taxonomic patterns in TDM. Other lower clades involved aerobic or facultative types like *Mesorhizobium* sp., *Saccharomonospora azure*a, *Ralstonia* sp. MD27, *Sporolactobacillus laevolacticus*. This work expands the community blueprints described by [4] that inhabit the Apatlaco river basin and suggest more diverse community patterns. Eventually, this type of comparative study between freshwater microbiomes and built environment metagenomes under xenobiotic nutritional selection will help to predict microbial shifts associated with anthropogenic activities.

### 3.2. Decolorization of Deep Blue 35 by the enriched microbiome

We measure the initial and final COD of the water composite sample during the enrichment process with Deep Blue 35, to appreciate the impact of the textile-dye community fraction on the overall organic load; which is a robust indicator of oxidizable pollutants found in water. The initial COD was 265.80 ± 5.0 mg.L^−1^ and the final COD was recorded as 133.33 ± 3.6 mg.L^−1^. Indicating that after 30 days on batch reactor the level of COD was reduced until values < 200 mg.L^−1^; in agreement with the Official Mexican Standard NOM-014-ECOL-1993, which establishes the permissible limits of pollutants in wastewater from the textile industry [36]. Also, we recorded the UV-VIS spectra of the dye-treated sample at the initial and final time (0-30 days by triplicated); the dye sample accounted for a discoloration rate of 23% (maximum visible wavelength: ~592-630 nm) (Figure 2). The absorption bands at ~630 nm - associated with the blue color-decreased slightly after 30 days on oligotrophic incubation, supporting that anthraquinone dye are both poorly adsorbed on to biomass or sediment. The relatively slight diminution of color essentially suggests that the removal of Deep Blue 35 was almost incomplete at this point, maybe due to a chromophore structure highly resistant to degradation, resulting in lengthy decolorization [37].

**Figure 2.**
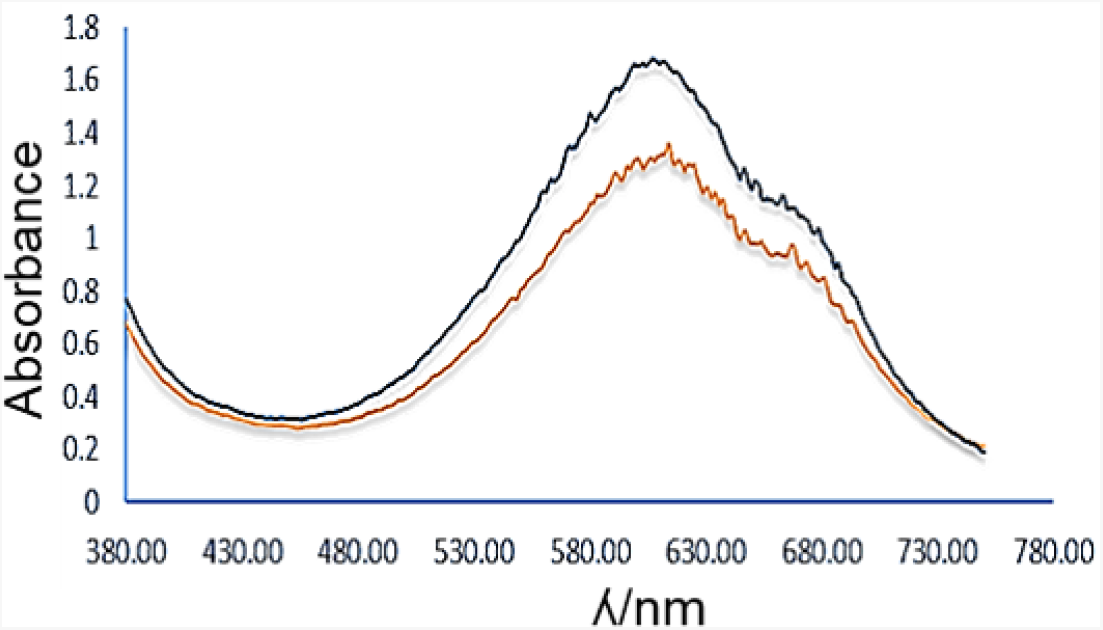
UV/Vis absorbance spectra recorded from the textile dye Deep-Blue 35(™) by the enriched microbiome. The blue line represents spectra recorded at day 0 with a concentration of 200 mg.L ^−1^ of textile dye. The yellow line represents spectra recorded after 30 days of incubation.

Although several strains capable of degrading anthraquinones dyes have been described, *Klebsiella* sp., *Pseudomonas* sp., *Bacillus cohnii* [38–40], very few studies prevail under oligotrophic conditions or oxygen limitation. *Bacillus cereus* DC11 has been reported as an anthraquinone degrader under oxygen limiting conditions, achieving a decolorization rate of Acid Blue 25 (100 μmol.L^−1^) near to 55% [41]. In this study we showed enrichment of anoxic and facultative taxa at the genus and species level (See taxonomic profile on Supplementary Material 1), suggesting that a reduced oxygen environment was created after Deep Blue 35 addition. Our results are numerically comparable to other achieved by single isolates like *Pseudomonas* sp. strain GM3 [42] with Reactive Blue 2 (degradation rate 14 %), *Aeromonas hydrophila* DN322 with Acid Blue 56 (degradation rate 21 %) [43]; *Enterobacter* sp. with Reactive Blue 19 (Degradation rate 25%) [44]. It is worthy to note that most of these cases were performed in rich media or supplemented with carbon sources such as glucose. Our approach roughly attempts to simulate what would happen in case of a discharge of textile effluent on to superficial waters and takes into account the modification of the microbiota as well as its ability to remove the contaminant in the absence of biostimulation. Altogether, suggest that the impact of textile dyes in freshwater could have long-term lasting effects before the indigenous microbiota been capable of modifying the pollutant significantly, this could represent conditions that promote anaerobiosis and methanogenesis. Otherwise, the microbial composition may experience significant modification.

### 3.3. Proximity-ligation metagenome deconvolution on textile dye selected microbiome

While MetaPhlAn provides useful taxonomic profiling through clade-specific markers, we attempt to obtain full genome community information by Hi-C metagenome deconvolution [8, 11]. ProxiMeta TDM deconvolution resulting in the reconstruction of 97 putative genome clusters (Figure 3a). Around 13.4% of the raw genome clusters have > 50% completeness; about 29.9% of the clusters have between 20-48% completeness, and 56.7% of genome composites have less than 20% completeness. Remarkably, 98% of the clusters produced by Hi-C have less than 10% of gene marker overrepresentation and 63 effective clusters possess less than 5% according to CheckM [19], supporting that Hi-C method produces clusters with little contamination levels [11, 12]. Forty-five clusters have a novelty score > 80% according to Mash distance metric [20], this anticipates the presence of an undescribed taxa representative from the Apatlaco basin (Supplementary Material 2); highlighting valuable novel genotaxonomic landscapes contained in the textile dye microbiome.

**Figure 3.**
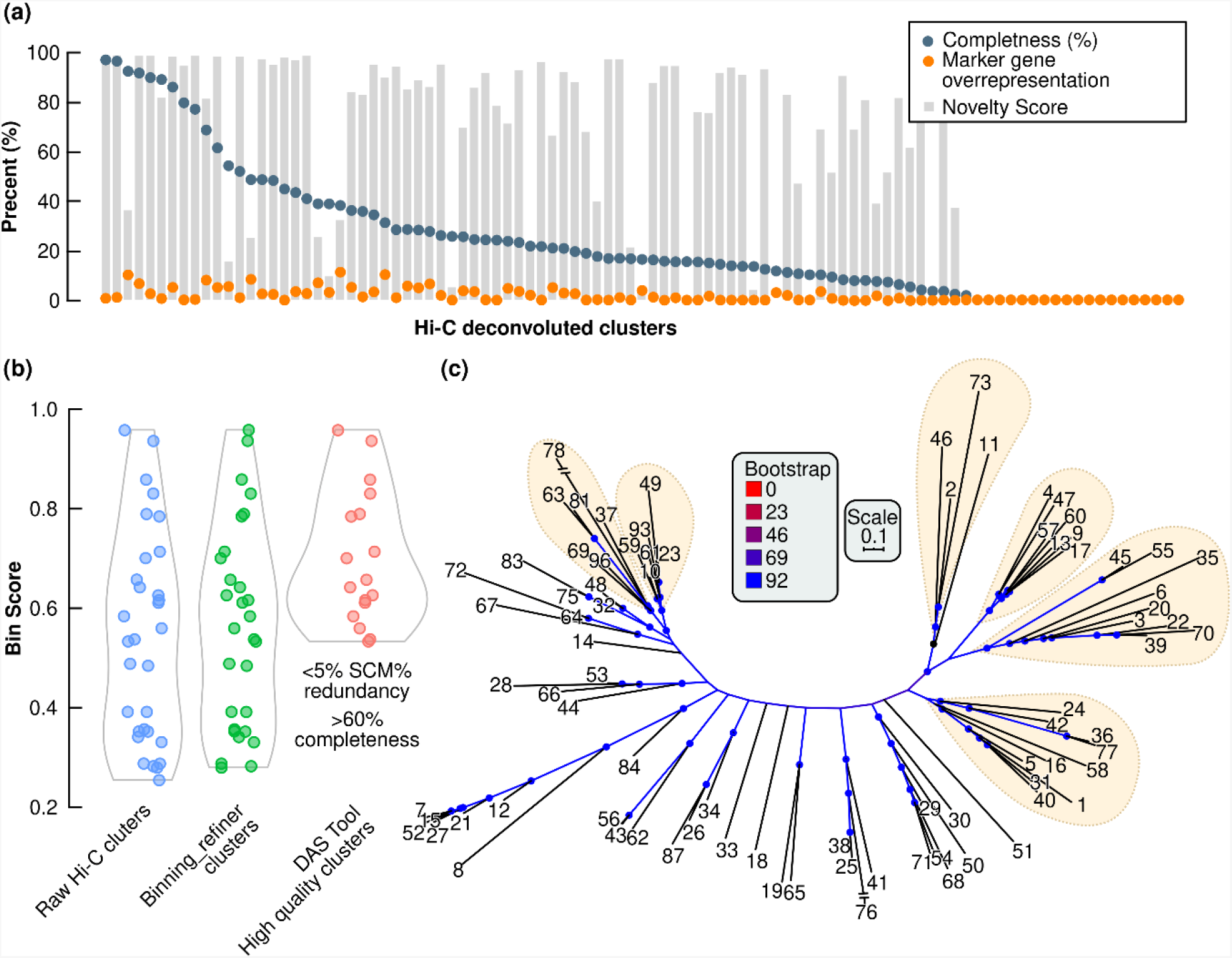
ProxiMeta Hi-C deconvoluted genomes in TDM. Quality criteria for genomes adhering to CheckM criteria for completeness (gray dots) and contamination (yellow dots), also the novelty score according to Mash D is showed (bars) (a). Refinement of Hi-C clusters to obtain optimized bins (b). Phylogenomic tree obtained with UBCG software grouping Hi-C clusters according to phylogenetic links (c); genome clusters with probable monophyletic relationship are enclosed in a light yellow leaf-like shape.

Subsequently, we applied Binner_refiner and DAS Tool overall 97 Hi-C assembly clusters to eliminate redundancy and generated more accurate genomes composites. This approach drastically reduced the total genome clusters to 18 improved drafts, all of them with medium or high-quality genomes properties [7, 21] (Figure 3B, Table 1). These 18 optimized metagenome-assembled genomes outperform the previous 14 Hi-C clusters with completeness ~50%. However, while we improved the overall quality of the most complete genomes, we do not rule out any of the Hi-C clusters obtained by proximity deconvolution for subsequent functional analysis; under the premise that these clusters came in principle from different cells in the microbiome, whose individual genomic complements may be functionally relevant. Full genome community exploration revealed novel species-level composites accurately; each Hi-C cluster represents an individual genomic unit (Intercluster Mash D > 0.2); this species segregation is properly reflected at phylogenetic level (Figure 3 C), with a clear differentiation between diverse taxonomic groups (*Archaea*, *Clostridiales*, *Actinobacteria*, *Proteobacteria*, or unknown taxa). This is a criterion to approximate species-specific relationships (two clusters of the same species share Mash D ≤ 0.05) in natural or build-environments microbiomes poorly-characterized. As in MetaPhlAn profiling, deconvolution produced clusters consistent with *Clostridales* and *Methanobacteriales* as the most abundant in the community. The combination of different tools for recovering metagenome-assembled genomes will improve our understanding of the structure and function of microbial communities in natural and synthetic ecosystems, revealing hidden species-level microbial contexts with his respective genomics complement.

**Table 1.** Optimized, non-redundant set of genome clusters from Hi-C textile-dye microbiome after trimming with DAS Tool and Binning refiner

| Cluster ID | Top Reference | Complete (%) | MGO (%) | Novelty score (%) | Abundance (%) | Conting N50 | Genome Size | Num Contings | GC (%) |
| --- | --- | --- | --- | --- | --- | --- | --- | --- | --- |
| Cluster 1* | *Clostridiales* | 98.10 | 1.02 | 98.36 | 2.28 | 96482 | 2,632,104 | 48 | 42.15 |
| Cluster 2* | Bacteria | 96.3 | 1.00 | 99.17 | 3.46 | 194975 | 5,263,022 | 43 | 61.00 |
| Cluster 3* | *Candidatus Afipia apatlaquensis* | 96.19 | 1.15 | 36.71 | 0.69 | 4194 | 5,604,749 | 1,705 | 60.72 |
| Cluster 4* | *Actinobacteria* | 91.79 | 1.10 | 99.13 | 0.83 | 3692 | 2,008,428 | 656 | 57.28 |
| Cluster 5* | *Clostridiales* | 94.29 | 1.06 | 99.11 | 1.11 | 16281 | 2,577,802 | 235 | 60.06 |
| Cluster 6* | *Acetobacter* sp. | 90.48 | 1.04 | 82.04 | 1.46 | 172292 | 2,397,974 | 29 | 60.31 |
| Cluster 7* | Euryarchaeota | 86.00 | 1.10 | 98.61 | 1.73 | 5192 | 2,230,951 | 608 | 38.87 |
| Cluster 8* | Euryarchaeota | 79.74 | 1.04 | 95.08 | 4.58 | 39230 | 2,697,761 | 86 | 36.82 |
| Cluster 9* | *Actinobacteria* | 87.62 | 1.02 | 98.97 | 1.41 | 26583 | 1,399,956 | 57 | 65.26 |
| Cluster 10* | *Clostridium* sp. | 64.76 | 1.07 | 81.64 | 6.22 | 12668 | 2,400,021 | 280 | 30.74 |
| Cluster 11* | Bacteria | 63.81 | 1.06 | 98.71 | 0.7 | 11943 | 3,932,167 | 488 | 52.21 |
| Cluster 12 | *Methanobacterium paludis* | 54.34 | 1.24 | 15.76 | 0.9 | 1696 | 2,501,351 | 1,461 | 35.96 |
| Cluster 13* | Actinobacteria | 59.05 | 1.05 | 98.48 | 1.11 | 12333 | 1,01,1957 | 100 | 67.89 |
| Cluster 14 | *Clostridium arbusti* | 48.74 | 1.34 | 25.22 | 1.49 | 4324 | 2,397,525 | 720 | 32.46 |
| Cluster 16* | *Oscillibacter* sp. | 67.62 | 1.13 | 95.12 | 0.91 | 3232 | 1,651,099 | 600 | 36.31 |
| Cluster 18* | *Clostridium* sp. | 59.05 | 1.10 | 97.27 | 1.73 | 4816 | 2,313,905 | 651 | 40.80 |
| Cluster 21 | *Methanobacterium paludis* | 48.37 | 3.20 | 10.02 | 1.10 | 1805 | 1,608,877 | 882 | 36.20 |
| Cluster 32 | *Clostridium arbusti* | 55.67 | 0.0 | 5.39 | 3.10 | 7238 | 683,706 | 126 | 30.08 |

### 3.4. Functional landscape selected by the textile dye, linking the phenotypic potential with individual genomes

To investigate the functional landscape distinctions between the textile dye and natural freshwater microbiomes, we concatenated all four freshwater metagenomes in a sole archive with KEGG annotations, to create a unified representative freshwater microbiome. Then we analyze the coding sequence elements related to major functional categories on KEGG reference database and identified relevant dye degradation enzymes coding sequences in both datasets.

Relatively similar amounts of coding sequences were assigned to textile and freshwater metagenomes (115,712 vs 135, 993; about ~ 10% difference between both predictions). Freshwater and textile dye metagenome fractions showed distinguishable functional profiles in the categories of carbohydrate, energy and lipid metabolism, glycan biosynthesis, xenobiotic biodegradation, and membrane transport. Interestingly, most of the molecular functions (KO) overrepresented in TDM, cluster independently (except for fructose and mannose metabolism) (Figure 4), suggesting a functional core significantly influenced by nutritional selection. Methane metabolism is a highly representative function in TDM probably enhanced after dye enrichment; this agrees with genomic observations where methanogenic taxa were the most abundant representatives (Supplementary Material 3). Also, we observed a higher frequency of *phnJ* gene (which encodes a releaser methane enzyme) in TDM (6 sequences elements) compared with natural microbiomes (1 sequence) [45]. Hence, it is expected that anaerobic mechanisms would be contending with the dye nutritional selection, insofar as other functional cores may be shaped in the community. Other functions like glycan synthesis and galactose metabolism could be related to cell wall synthesis processes and nucleotide sugar biosynthesis to support active growth. The phosphotransferase system could be triggered in response to oligotrophic conditions, to potentiate uptake of carbohydrates from a medium limited in nutrients [46]. Likewise, in the freshwater fraction, the most representative metabolic pathway was related to the ABC transporters system, followed by the fatty acid, aromatic degradation, and lipopolysaccharide biosynthesis. This is consistent with the reported by Breton-Deval *et al*., [27] and support functions associated with the metabolism of anthropogenic substances coming from diverse origins (industrial, agricultural, and domestic discharges).

**Figure 4.**
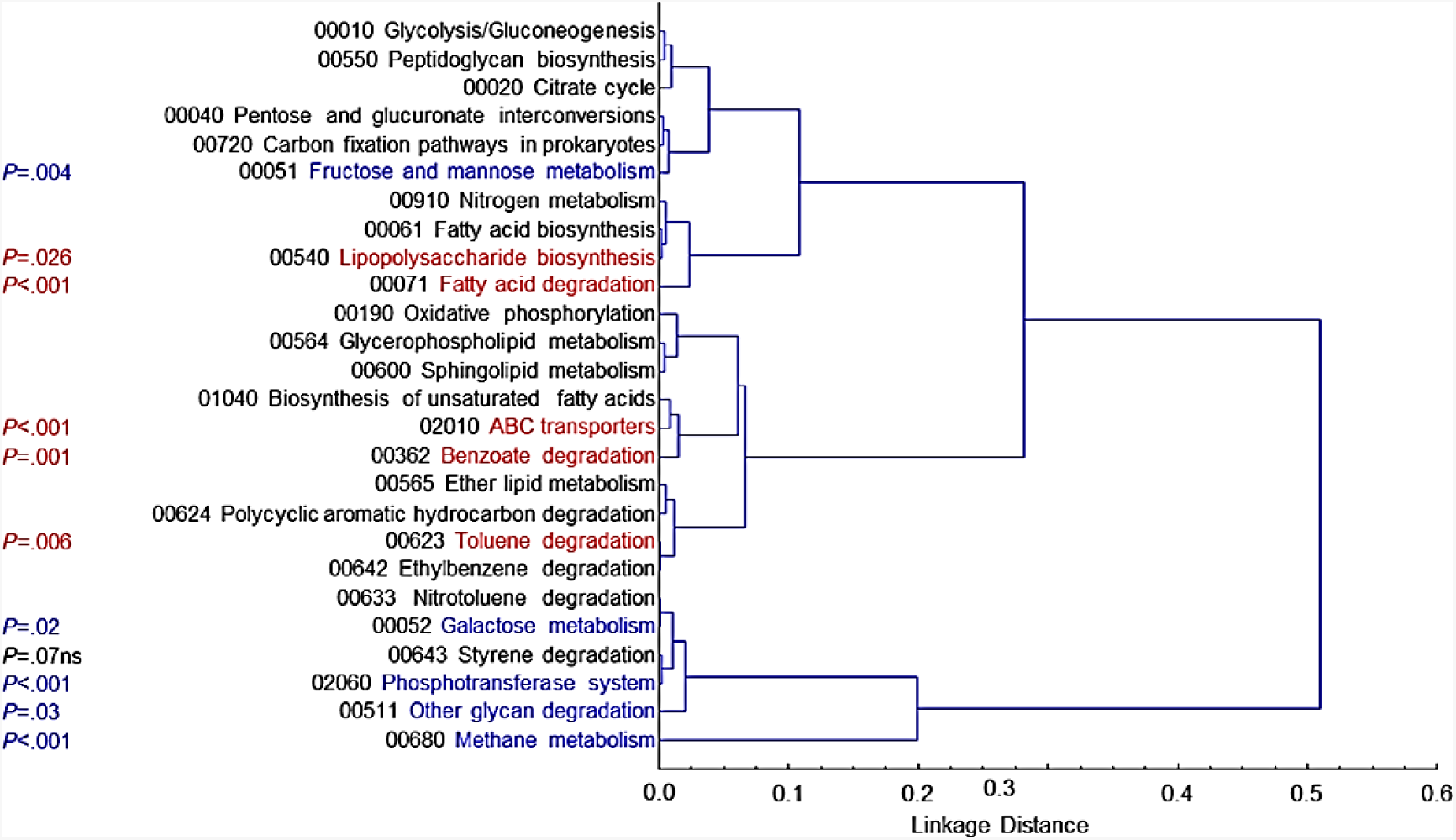
Main KEGG metabolic pathways in the TDM compared to natural freshwater metagenome. Overrepresented pathways are depicted in blue for TDM and red for the freshwater metagenome. P values are depicted on the left side.

Subsequently, we constructed a presence-absence matrix with dye-decoloring enzymes deduced from KEGG annotations in TDM (see MM), and clustered against the genome composites they came from; under the premise that they constitute potential phenotypic markers associable with the ability to degrade textile dyes (Table 2).

**Table 2.** Coding sequence abundance of dye-decoloring related enzymes in freshwater and textile dye microbiomes according to KEGG annotations

| KO | Freshwater microbiome | Textile dye microbiome | EC Number | Name | Gene name |
| --- | --- | --- | --- | --- | --- |
| K15733 | 2 | - | 1.11.1.19 | Dye decolorizing peroxidase | *DyP* |
| K03206 | 1 | 1 | 1.7.1.6 | Azoreductase | *azr* |
| K01118 | 11 | 26 | 1.7.1.17 | NADH-azoreductase | *acpD, azoR* |
| K05909 | 1 | 1 | 1.10.3.2 | Laccase | - |
| K03782 | 142 | 13 | 1.11.1.21 | Catalase-peroxidase | *katG* |
| K05810 | 42 | 21 | 1.10.3.- | Polyphenol oxidase | - |

Dye biodegradation trait could be connected efficiently to emerging new taxa deconvoluted in TDM based on dye biodegradation functional genes (Figure 5). In agreement with the overall distribution, NADH-Azoreductase, Polyphenol oxidase, and Catalase-peroxidase sequence elements were the most abundant per genome. Functional redundancy was observed mainly on the clusters 1 (novel taxon); 3 (*Candidatus* Afipia apatlaquensis); 7 (novel *Euryarchaeota* member); 11 (novel taxon); 14 (*Clostridium arbusti*); 20 (*Mesorhizobium* sp., novel species) and 41 (*Cupriavidus metallidurans*) with three or more copies by genome. *Clostridium* and *Mesorhizobium* members have been previously identified as capable to degrade anthraquinonic dyes [47][48], *C. metallidurans* is largely known for his tolerance to metals and have been reported in aromatic compound degradation like toluene [49].

**Figure 5.**
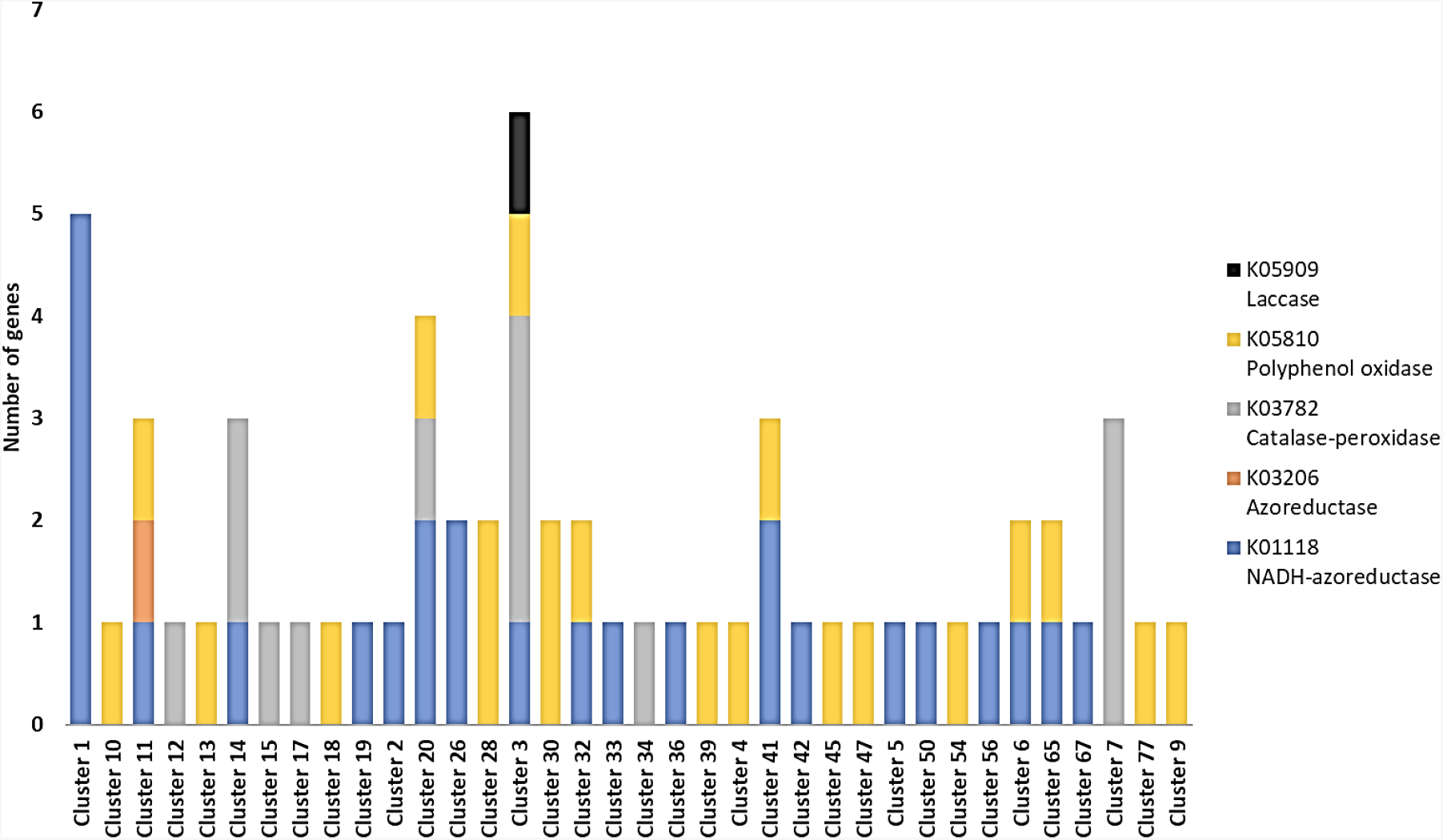
Abundance of dye-decolorizing coding sequences in the TDM deconvoluted genomes. The enzymes selected are color labeled and his KO is shown on the right side.

#### 3.4.1 Sequence elements significantly associated with simulated phenotype based on genotype

To support the hypothesis that it is possible to connect individual genomic composites with explicit functions within a community under nutritional selection, we apply an association analysis of sequence elements with phenotypes through a genome-wide association approach. We artificially associated a binary predicted phenotype to each genome composites from this study, based on the former presence/absence of dye-decoloring coding sequences. By applying Seer tool [29] over the 97 TDM genome composites we predicted 17, 194,445 genetics features (total k-mers). After data filtration 57,115 final k-mers were printed as putative elements with effect on textile dye biodegradation phenotype, and 43,141 significant k-mers aligned with sequence elements related to biodegradation of textile dyes. The genetic determinants mostly associated with the biodegradation of textile dyes were those corresponding to Catalase-peroxidase > Polyphenol oxidase > Laccase (Table 3). The clusters 3, 14 (*Clostridium arbusti*), and 41(*Cupriavidus metallidurans*) also remark as attractive candidates for targeted studies on the degradation of textile dyes. The cluster 3, (identified as “*Candidatus* Afipia apatlaquensis ”) genome composite has been recently suggested as a tentative textile dye biodegrader [50]. Also totally novel taxa from different fila (*Euryarchaeota*, *Firmicutes*, *Proteobacteria*) are significantly associated with the textile-dye degradation phenotype, expanding the catalog of microbiodiversity related to textile biodegradation. This suggests that these sequence elements are strongly associated with the predicted phenotype, compared to those with lower coverage, and tentatively could have a greater effect on it.

**Table 3.** Metagenome association results for the textile dye biodegradation predicted phenotype over 97 genome composites on the TDM community. Predicted phenotype was based on dye-decoloring coding sequence presence/absence. Significant k-mers were mapped to Prokka annotations for all dye-decoloring related genes in the community. The hits with significant k-mers > 1000 mapped to reference are shown

| Genome representation | KO | Functional annotation or Conserved domains | k-mers mapped | Bases with coverage (%) | Average coverage depth |
| --- | --- | --- | --- | --- | --- |
| Cluster 7 (*Methanobacterium* sp.)* | K03782 | Catalase-peroxidase | 2117 | 99.35 | 9.76 |
| Cluster 17 (*Proteobacteria*)* | K03782 | Catalase-peroxidase | 1689 | 99.65 | 7.77 |
| Cluster 34 (*Clostridium* sp.)* | K03782 | Catalase-peroxidase | 1678 | 98.29 | 8.19 |
| Cluster 14 (*Clostridium arbusti*) | K03782 | Catalase-peroxidase | 1404** | 100 | 12.47 |
|  |  |  | 1237** | 99.81 | 11.78 |
| Cluster 12 (*Methanobacterium paludis*) | K03782 | Catalase-peroxidase | 1261 | 99.08 | 8.92 |
| Cluster 3 (*Candidatus Afipia apatlaquensis*) | K03782 | Catalase-peroxidase | 1065 | 98.39 | 7.46 |
|  | K05909 | Laccase | 1001 | 97.42 | 7.05 |
| Cluster 41 (*Cupriavidus metallidurans*) | K05810 | Polyphenol oxidase | 1050 | 100 | 13.09 |

\*Novel species: Mash D > 0.05 with it closets neighbor
\*\*k-mers mapped to two different sequence elements with common orthology

As genome encodes the metabolic abilities of microorganisms it is a major genetic causal of phenotypic behaviors. However, the prediction of individual organismal functions directly from genes must be taken with caution. In this way, we propose that the genomes reported in figure 5 could be involved in the degradation of textile dyes based on the genetic potential they show. This would be useful on rapid screenings within a large collection of genomes in order to select the most represented functional feature (for example in the dissection of genome composites with the greatest contribution to an observed or hypothetical phenotype), reducing costs and experimental efforts. As well as predicting phenotypes within complex communities under environmental pressures.

## Conclusions

These results help to understand how nutritional selection by a xenobiotic substance (anthraquinone textile dye) shapes local microbial landscapes at the community level. Over 50% of the total microbial diversity is related to anoxygenic lifestyle, also methanogenic functions were significantly enriched in the textile microbiome, supporting major role for anaerobic contexts in the microbiome. Organic oxidizable content was reduced by 50% with a modest decolorization rate, suggesting that more time is needed to achieve complete removal of complex textile dye structure. We postulate that sequence elements related to catalase-peroxidase, polyphenol oxidase, and laccase enzymes may be causally associated with textile-dye degradation phenotypes and may be putative markers for environmental monitoring of decolorizing functions, as in textile wastewater treatment plants. This study prompts rapid genomic screening in order to select statistically represented functional features, reducing costs, and experimental efforts. As well as predicting phenotypes within complex communities under environmental pressures.

## Acknowledgments

The authors thanks UNAM-IBT, for support this study through Project 2030 P-10065. ASR thank to the program CATEDRAS CONACYT from the Consejo Nacional de Ciencia y Tecnología, Mexico, for support the Project 237. Hayley Mangelson was supported in part by NIH grants 5R44AI122654 and 1R44AI150008 to Phase Genomics. We also like to thank the Unidad Universitaria de Secuenciación Masiva y Bioinformática (UUSMB) of the Instituto de Biotecnología, UNAM and Dr. Miguel Lara Flores for their support.

## References

1. Parks DH, Rinke C, Chuvochina M, et al (2017) Recovery of nearly 8,000 metagenome-assembled genomes substantially expands the tree of life. Nat Microbiol. https://doi.org/10.1038/s41564-017-0012-7

2. Yang P, Su X, Ou-Yang L, et al (2014) Microbial community pattern detection in human body habitats via ensemble clustering framework. BMC Syst Biol 8:S7. https://doi.org/10.1186/1752-0509-8-s4-s7

3. Alneberg J, Bennke C, Beier S, et al (2020) Ecosystem-wide metagenomic binning enables prediction of ecological niches from genomes. Commun Biol 3:. https://doi.org/10.1038/s42003-020-0856-x

4. Breton-Deval L, Sanchez-Flores A, Juárez K, Vera-Estrella R (2019) Integrative study of microbial community dynamics and water quality along The Apatlaco River. Environ Pollut. https://doi.org/10.1016/j.envpol.2019.113158

5. Kang DD, Froula J, Egan R, Wang Z (2015) MetaBAT, an efficient tool for accurately reconstructing single genomes from complex microbial communities. PeerJ 3:e1165. https://doi.org/10.7717/peerj.1165

6. Albertsen M, Hugenholtz P, Skarshewski A, et al (2013) Genome sequences of rare, uncultured bacteria obtained by differential coverage binning of multiple metagenomes. Nat Biotechnol 31:533–538. https://doi.org/10.1038/nbt.2579

7. Sieber CMKK, Probst AJ, Sharrar A, et al (2018) Recovery of genomes from metagenomes via a dereplication, aggregation and scoring strategy. Nat Microbiol 3:836–843. https://doi.org/10.1038/s41564-018-0171-1

8. Burton JN, Liachko I, Dunham MJ, Shendure J (2014) Species-level deconvolution of metagenome assemblies with Hi-C-based contact probability maps. G3 (Bethesda) 4:1339–46. https://doi.org/10.1534/g3.114.011825

9. Marbouty M, Koszul R (2015) Metagenome Analysis Exploiting High-Throughput Chromosome Conformation Capture (3C) Data. 31:673–682. https://doi.org/10.1016/j.tig.2015.10.003

10. Stewart RD, Auffret MD, Warr A, et al (2018) Assembly of 913 microbial genomes from metagenomic sequencing of the cow rumen. Nat Commun 9:870. https://doi.org/10.1038/s41467-018-03317-6

11. Press MO, Wiser AH, Kronenberg ZN, et al (2017) Hi-C deconvolution of a human gut microbiome yields high-quality draft genomes and reveals plasmid-genome interactions. bioRxiv. https://doi.org/10.1101/198713

12. Gaytán I, Sánchez-Reyes A, Burelo M, et al (2020) Degradation of Recalcitrant Polyurethane and Xenobiotic Additives by a Selected Landfill Microbial Community and Its Biodegradative Potential Revealed by Proximity Ligation-Based Metagenomic Analysis. Front Microbiol 10:. https://doi.org/10.3389/fmicb.2019.02986

13. Lieberman-Aiden E, Berkum N van (2009) Comprehensive mapping of long range interactions reveals folding principles of the human genome. Science (80-) 326:289–293. https://doi.org/10.1126/science.1181369.Comprehensive

14. Bushnell B, Rood J, Singer E (2017) BBMerge – Accurate paired shotgun read merging via overlap. PLoS One. https://doi.org/10.1371/journal.pone.0185056

15. Li D, Luo R, Liu CM, et al (2016) MEGAHIT v1.0: A fast and scalable metagenome assembler driven by advanced methodologies and community practices. Methods

16. Li H, Durbin R (2009) Fast and accurate short read alignment with Burrows-Wheeler transform. Bioinformatics. https://doi.org/10.1093/bioinformatics/btp324

17. Faust GG, Hall IM (2014) SAMBLASTER: Fast duplicate marking and structural variant read extraction. In: Bioinformatics

18. Li H, Handsaker B, Wysoker A, et al (2009) The Sequence Alignment/Map format and SAMtools. Bioinformatics. https://doi.org/10.1093/bioinformatics/btp352

19. Parks DH, Imelfort M, Skennerton CT, et al (2015) CheckM: assessing the quality of microbial genomes recovered from. Genome Res 25:1043–1055. https://doi.org/10.1101/gr.186072.114

20. Ondov BD, Treangen TJ, Melsted P, et al (2016) Mash: Fast genome and metagenome distance estimation using MinHash. Genome Biol 17:1–14. https://doi.org/10.1186/s13059-016-0997-x

21. Song WZ, Thomas T (2017) Binning-refiner: Improving genome bins through the combination of different binning programs. Bioinformatics. https://doi.org/10.1093/bioinformatics/btx086

22. Truong DT, Franzosa EA, Tickle TL, et al (2015) MetaPhlAn2 for enhanced metagenomic taxonomic profiling. Nat. Methods

23. Asnicar F, Weingart G, Tickle TL, et al (2015) Compact graphical representation of phylogenetic data and metadata with GraPhlAn. PeerJ. https://doi.org/10.7717/peerj.1029

24. Seemann T (2014) Prokka: Rapid prokaryotic genome annotation. Bioinformatics. https://doi.org/10.1093/bioinformatics/btu153

25. Aramaki T, Blanc-Mathieu R, Endo H, et al (2019) KofamKOALA: KEGG ortholog assignment based on profile HMM and adaptive score threshold. Bioinformatics. https://doi.org/10.1093/bioinformatics/btz859

26. Sarkar S, Banerjee A, Halder U, et al (2017) Degradation of Synthetic Azo Dyes of Textile Industry: a Sustainable Approach Using Microbial Enzymes. Water Conserv Sci Eng. https://doi.org/10.1007/s41101-017-0031-5

27. Breton-Deval L, Sanchez-Reyes A, Sanchez-Flores A, et al (2020) Functional Analysis of a Polluted River Microbiome Reveals a Metabolic Potential for Bioremediation. Microorganisms 8:554. https://doi.org/10.3390/microorganisms8040554

28. Kanehisa M, Goto S, Sato Y, et al (2012) KEGG for integration and interpretation of large-scale molecular data sets. Nucleic Acids Res. https://doi.org/10.1093/nar/gkr988

29. Lees JA, Vehkala M, Välimäki N, et al (2016) Sequence element enrichment analysis to determine the genetic basis of bacterial phenotypes. Nat Commun. https://doi.org/10.1038/ncomms12797

30. Seemann T (2018) barrnap 0.9 : rapid ribosomal RNA prediction

31. Na SI, Kim YO, Yoon SH, et al (2018) UBCG: Up-to-date bacterial core gene set and pipeline for phylogenomic tree reconstruction. J Microbiol 56:281–285. https://doi.org/10.1007/s12275-018-8014-6

32. Barco RA, Garrity GM, Scott JJ, et al (2020) A genus definition for bacteria and archaea based on a standard genome relatedness index. MBio. https://doi.org/10.1128/MBIO.02475-19

33. Satola B, Wübbeler JH, Steinbüchel A (2013) Metabolic characteristics of the species Variovorax paradoxus. Appl. Microbiol. Biotechnol.

34. Öztürk B, Werner J, Meier-Kolthoff JP, et al (2020) Comparative genomics suggests mechanisms of genetic adaptation towards the catabolism of the phenylurea herbicide linuron in Variovorax. Genome Biol Evol. https://doi.org/10.1093/gbe/evaa085

35. Dréno B, Pécastaings S, Corvec S, et al (2018) Cutibacterium acnes (Propionibacterium acnes) and acne vulgaris: a brief look at the latest updates. J. Eur. Acad. Dermatology Venereol.

36. NOM-CCA-014-ECOL-1993 (1993) NORMA Oficial Mexicana NOM-CCA-014-ECOL/1993, que establece los límites máximos permisibles de contaminantes en las descargas de aguas residuales a cuerpos receptores provenientes de la industria textil. México

37. Routoula E, Patwardhan S V. (2020) Degradation of Anthraquinone Dyes from Effluents: A Review Focusing on Enzymatic Dye Degradation with Industrial Potential. Environ Sci Technol 54:647–664. https://doi.org/10.1021/acs.est.9b03737

38. Xie X, Liu N, Yang B, et al (2016) Comparison of microbial community in hydrolysis acidification reactor depending on different structure dyes by Illumina MiSeq sequencing. Int Biodeterior Biodegrad. https://doi.org/10.1016/j.ibiod.2016.04.004

39. Forss J, Lindh M V., Pinhassi J, Welander U (2017) Microbial biotreatment of actual textile wastewater in a continuous sequential rice husk biofilter and the microbial community involved. PLoS One. https://doi.org/10.1371/journal.pone.0170562

40. Mishra S, Maiti A (2018) The efficacy of bacterial species to decolourise reactive azo, anthroquinone and triphenylmethane dyes from wastewater: a review. Environ. Sci. Pollut. Res.

41. Deng D, Guo J, Zeng G, Sun G (2008) Decolorization of anthraquinone, triphenylmethane and azo dyes by a new isolated Bacillus cereus strain DC11. Int Biodeterior Biodegrad. https://doi.org/10.1016/j.ibiod.2008.01.017

42. Yu J, Wang X, Yue PL (2001) Optimal decolorization and kinetic modeling of synthetic dyes by pseudomonas strains. Water Res. https://doi.org/10.1016/S0043-1354(01)00100-2

43. Ren S, Guo J, Zeng G, Sun G (2006) Decolorization of triphenylmethane, azo, and anthraquinone dyes by a newly isolated Aeromonas hydrophila strain. Appl Microbiol Biotechnol. https://doi.org/10.1007/s00253-006-0418-2

44. Holkar CR, Pandit AB, Pinjari D V. (2014) Kinetics of biological decolorisation of anthraquinone based Reactive Blue 19 using an isolated strain of Enterobacter sp.F NCIM 5545. Bioresour Technol. https://doi.org/10.1016/j.biortech.2014.09.108

45. Yao M, Henny C, Maresca JA (2016) Freshwater bacteria release methane as a by-product of phosphorus acquisition. Appl Environ Microbiol. https://doi.org/10.1128/AEM.02399-16

46. Biddanda B, Ogdahl M, Cotner J (2001) Dominance of bacterial metabolism in oligotrophic relative to eutrophic waters. Limnol Oceanogr. https://doi.org/10.4319/lo.2001.46.3.0730

47. Li H hong, Wang Y tao, Wang Y, et al (2019) Bacterial degradation of anthraquinone dyes. J. Zhejiang Univ. Sci. B

48. Lalnunhlimi S, Veenagayathri K (2016) Decolorization of azo dyes (Direct Blue 151 and Direct Red 31) by moderately alkaliphilic bacterial consortium. Brazilian J Microbiol. https://doi.org/10.1016/j.bjm.2015.11.013

49. Tofalos AE, Daghio M, González M, et al (2018) Toluene degradation by Cupriavidus metallidurans CH34 in nitrate-reducing conditions and in Bioelectrochemical Systems. FEMS Microbiol. Lett.

50. Sánchez-Reyes A, Bretón-Deval L, Mangelson H, Sanchez-Flores A (2020) Draft genome sequence of “ Candidatus Afipia apatlaquensis ” sp. nov., IBT ‑ C3, a potential strain for decolorization of textile dyes. BMC Res Notes 13:1–3. https://doi.org/10.1186/s13104-020-05117-y

